# MoTERNN: Classifying the Mode of Cancer Evolution Using Recursive Neural Networks

**DOI:** 10.1101/2022.08.21.504710

**Authors:** Mohammadamin Edrisi, Huw A. Ogilvie, Meng Li, Luay Nakhleh

## Abstract

With the advent of single-cell DNA sequencing, it is now possible to infer the evolutionary history of thousands of tumor cells obtained from a single patient. This evolutionary history, which takes the shape of a tree, reveals the mode of evolution of the specific cancer under study and, in turn, helps with clinical diagnosis, prognosis, and therapeutic treatment. In this study we focus on the question of determining the mode of evolution of tumor cells from their inferred evolutionary history. In particular, we employ recursive neural networks that capture tree structures to classify the evolutionary history of tumor cells into one of four modes—*linear*, *branching*, *neutral*, and *punctuated*. We trained our model, MoTERNN, using simulated data in a supervised fashion and applied it to a real phylogenetic tree obtained from single-cell DNA sequencing data. MoTERNN is implemented in Python and is publicly available at https://github.com/NakhlehLab/MoTERNN.

## 1 Introduction

From an evolutionary perspective, clonal evolution in cancer and intra-tumor heterogeneity (ITH) are the results of an interplay between mutations and selective pressures in the tumor micro-environment [1–3] and can be in part some of the contributing factors in metastasis [4] and tumor drug resistance [5, 6]. Aided by advances in sequencing technologies such as microarray [7], next-generation sequencing [8, 9], and single-cell sequencing [10, 11] that have been developed over the last three decades, the field of cancer evolution has gained attention as studies have shown supporting evidence of tumor cells being subject to selective pressures in response to their environment, which includes the immune system response as well as treatments such as chemotherapy and radiation.

As cancer cells are sampled and sequenced at a small number of time points (most often only one), understanding cancer evolution is done by inferring the evolutionary history of the sampled cells from their somatic mutations—single-nucleotide variations (SNVs) and copy number aberrations (CNAs)—obtained by a variety of DNA sequencing technologies. Indeed, a wide array of cancer evolutionary tree inference methods has been developed in the last decade, which use bulk sequencing data, single-cell sequencing data or a combination thereof [12–27].

While evolution is a stochastic process, it has been shown that this process could be constrained in cancer as evidenced by different modes of evolution observed in different cancers and, sometimes, during the lifetime of the same cancer (e.g., see Table 1 in [28]). The four main modes of evolution are linear evolution (LE), branching evolution (BE), neutral evolution (NE), and punctuated evolution (PE), all of which are illustrated in Fig.1. In linear evolution, some cells acquire somatic mutations with strong selective advantages over other cells. This selective sweep results in the tumor being dominated by a major *clone*^1^ and a few persistent minor clones that survived from the previous selective sweeps. Thus, the expected phylogenetic tree would take a ladder-like shape, as illustrated in Fig. 1a. In branching evolution, the clones evolve in parallel while all gaining fitness during their evolution. Consequently, multiple clones are expected to be present at the time of tissue sampling [29], and the phylogenetic tree would take an overall balanced shape as the clones do not outcompete each other. However, in each clone, one can observe fitness changes during the lifetime of the tumor (Fig. 1b). Neutral evolution refers to the case where there is no selection during the lifespan of the tumor. Neutral evolution assumes that the accumulation of mutations is merely a result of tumor progression and natural selection does not play much of a role; thus, it provides an alternative explanation for patterns and frequencies of mutations [29,30]. The expected shape of a phylogeny following this model would be highly balanced, not only on the tree’s backbone but also at the level of individual cells, as illustrated in Fig. 1c. Finally, punctuated evolution, first proposed in paleontology [31], is based on the idea that tumor progression begins in a *big bang* fashion: at the earlier stages, there is a burst of a large number of mutations. Following this phase, the clones gradually grow, leading to a few dominant clones in the tumor. Since the burst of mutations occurs earlier, all the clones share a large portion of mutations yielding a long root branch in the expected phylogeny, as illustrated in Fig. 1d.

**Figure 1:**
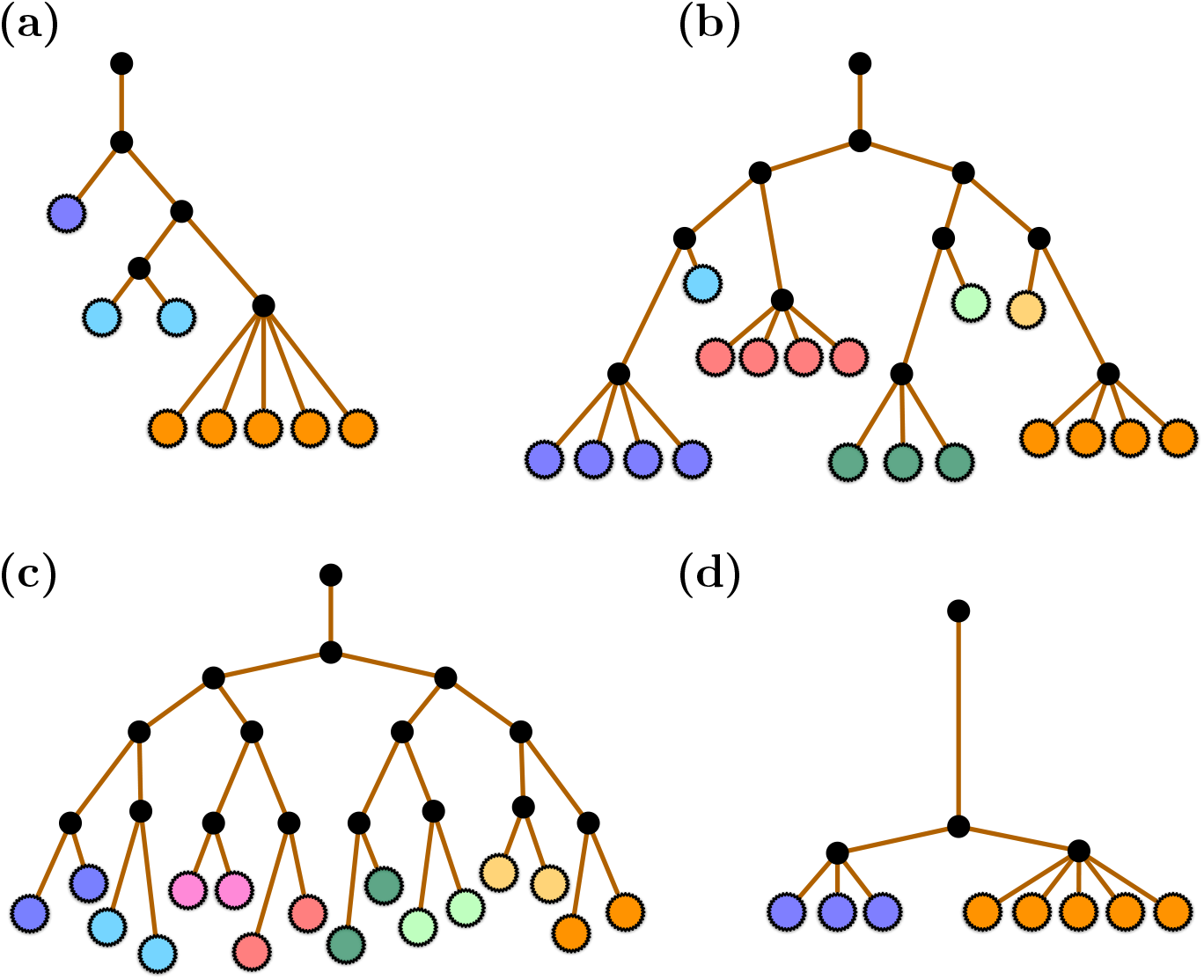
The phylogenetic trees indicative of four different modes of evolution. (a) Linear evolution, where a clone takes over the cancer cell population. (b) Branching evolution, where multiple clones arise and evolve in parallel at different rates due to selective pressures. (c) Neutral evolution, where multiple clones arise and evolve at similar rates. (d) Punctuated evolution, where a burst of mutations occurs followed by the growth of clones. Ancestral cells are shown in black solid circles, whereas present-day cells are shown in colored solid circles. (Reproduced from [29])

Determining the mode of tumor evolution is important as these models have different diagnostic, prognostic, and therapeutic implications [28]. For example, a tumor following LE and PE models would require a simpler biopsy since a few samples are good representatives of the entire tumor. On the other hand, BE and NE indicate a high degree of ITH and thus require more biopsy samples for diagnostic purposes [29].

Common approaches to determining the mode of cancer evolution include simulations and mathematical modeling. There is a rich body of literature on mathematical modeling of tumor evolution based on stochastic processes such as the multi-type branching evolution process [32] and the Moran process [33]. These stochastic processes provide predictive statistic measurements on the population size of cancer cells, mutant allele frequencies, or mutation rates [32,34] whose agreement with the observed data determines the mode of evolution. As an example of a simulation-based approach, [35] identified punctuated evolution in triple-negative breast cancer (TNBC) by simulating CNA phylogenetic trees under gradual and punctuated modes of multi-type stochastic birth-death-mutation process, and then measured the fitness of each simulation scenario to the real data using AMOVA analysis [36]. Although this approach benefits from taking the evolutionary history of cells into account, generating realistic phylogenies is still one of the challenges in the field. [37] assessed the fitness of two evolutionary hypotheses to eight TNBC tumors. In both models, the tumor growth starts with a punctuated burst of CNA events. In one model this punctuated phase is followed by a “*gradual accumulation of CNAs*” at a constant rate, and in the other model it leads to a “*transient instability*” in genomic evolution, then a return to the gradual evolution phase. Incorporating these two models into a likelihood framework enabled the authors to measure the fitness of the two models using Akaike Information Criterion [38]. This analysis showed punctuated evolution followed by transient instability and gradual evolution better describes the TNBC tumors. In addition to these approaches, one can use model-based approaches (e.g., in [39]) that accurately detect the speciation events that agree with the PE mode of evolution.

Outside the above categories, Phyolin [40] and the method of [41] identify the mode of evolution given the binary genotype matrices obtained from single-cell SNVs. These methods, however, are aimed at distinguishing between linear and nonlinear modes (i.e., a binary classification), which is a simpler problem than the one we address here.

In this study, we tackle the problem of determining the mode of cancer evolution as a graph classification where the graphs are phylogenetic trees and their labels/classes are the corresponding evolutionary modes. Specifically, we investigate the appropriateness of recursive neural networks, or RNNs^2^, first proposed in [42]. RNNs have been successfully used in natural language processing to capture semantic relationships between words in variable-sized sentences and parse trees [42, 43]. Here, the *words* and their *semantic representations* are the genotypes of the present-day cells and their ancestors, respectively. It is worth mentioning that the capability of RNNs in exploiting the tree structure of phylogenies makes them a more natural candidate for our task than stand-alone multi layer perceptrons (MLPs), convolutional neural networks (CNNs), or traditional classification methods such as random forests. To train our model in a supervised manner, we modified the beta-splitting model (BSM) [44] to generate simulated phylogenetic trees and genotypes according to the four cancer evolutionary modes. Next, we applied our model, MoTERNN (<u>Mo</u>de of <u>T</u>umor <u>E</u>volution using <u>R</u>ecursive <u>N</u>eural <u>N</u>etworks), on a real biological phylogenetic tree obtained from single-cell DNA sequencing (scDNAseq) data of a TNBC patient [45]. MoTERNN classified the mode of evolution as punctuated evolution, in agreement with the original study, even though we used SNV data here while the original study used CNA data.

Our study demonstrates the suitability of RNNs for classification problems on tree structures. By developing MoTERNN, we have added to the evolutionary biology toolbox that is used to study and understand cancer biology.

## 2 Methods

### 2.1 Problem description

The input to MoTERNN consists of a genotype matrix and a phylogenetic tree obtained from tumor samples. Let **G** = (*g_ij_*) ∈ {0, 1}^*N*×*M*^ be a binary genotype matrix where 0 and 1 indicate absence and presence of mutations, respectively, *N* is the number of samples, and *M* is the number of genomic loci. Following this notation, *g_i_* represents the genotype vector of the *i*^th^ sample at the *j*^th^ locus. Let 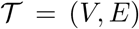 be a phylogenetic tree where *V* and *E* are the sets of nodes and edges, respectively. In this work, we assume that the phylogenetic tree is binary. The leaves of 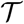 are bijectively labeled by the genotype vectors (rows) in **G**. Given **G** and 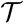, MoTERNN predicts one of the four labels in the set {LE, BE, NE, PE}. Next, we give a brief background on RNNs and then describe the model underlying MoTERNN.

### 2.2 Recursive Neural Networks

Dating back to the late eighties and early nineties, training neural networks on recursive data structures attracted interest from the machine learning community. Seminal works on models such as recursive autoassociative memory [46] and backpropagation through structure (BTS) [47] established the basis for dealing with variable-sized recursive structures and efficient computation of backpropagation. Utilizing the BTS scheme for backpropagation, [42] introduced RNNs for parsing natural scene images and natural language sentences.

The input of an RNN is a binary tree (or the corresponding adjacency matrix) and a set of vector embeddings for all the leaves. The goal is to learn and predict the labels of the internal nodes including the root of the tree. To predict the label of a node, a corresponding vector embedding for the node is required. In our case, the embeddings^3^ of the leaves are generated using an encoder neural network that maps the genotype profiles into a shared lower dimension. Given a tree, the algorithm traverses it and computes the labels and embeddings of the internal nodes recursively. The embedding of a parent node is computed by a neural network which takes as input the concatenation of the two children’s embeddings and generates the parent’s embedding as output. Following the terminology from [43], we refer to this network as a *compositionality function*. To predict a node’s label, its embedding is given to a classifier neural network that produces scores for the node’s association to each class. It is worth mentioning that what separates RNNs from their predecessors is that the same compositionality function and classifier network are applied to all inputs resulting in a more flexible and computationally efficient architecture.

Although the original RNN aimed at predicting the labels of all nodes, we associate the evolutionary model of a phylogenetic tree with its root. Thus, in our scheme, the other internal nodes do not have labels, and the RNN only predicts the root’s label. Next, we describe the three components of our model including the encoder, compositionality function, and classifier.

### 2.3 Encoder network

Each genotype profile *g_i_* is mapped into a shared lower dimension using an encoder network *ϕ*:

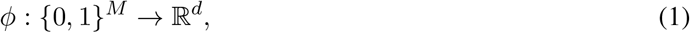

where *d* is a user-specified parameter. Although the encoder network *ϕ* is applied to only the genotypes at the leaves of 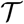, we use *ϕ*(*v*) more generally to denote the embedding of any node *v* in the tree 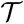. In our implementation, we used a single-layer feed-forward neural network with *d* = 128 as our encoder network.

### 2.4 Compositionality function

The compositionality function is a neural network that computes the embedding of a parent node given the computed embeddings of its children. Let *v* be an internal node and *ϕ*(*v*_1_) and *ϕ*(*v*_2_) be the embeddings of its children *v*_1_ and *v*_2_. The embedding of *v* based on the compositionality function is computed using the following formula:

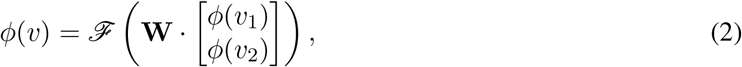

where 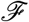 is a non-linear activation function (such as ReLU) and 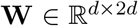 represents the weights of the compositionality function which are multiplied by the concatenation of the two embeddings of the children. Fig. 2a illustrates an example of a tree being traversed by an RNN and how the compositionality function operates recursively during this process. First, the embeddings of the leaves *A, B*, and *C* are obtained from the encoder network. The leaves’ embeddings are denoted by *ϕ*(*A*), *ϕ*(*B*), and *ϕ*(*C*). The first internal node whose embedding is computed is *D* because its children’s embeddings have already been computed. Next, the embedding of the root node, *E*, is computed by passing *ϕ*(*C*) and *ϕ*(*D*) to the compositionality function.

**Figure 2:**
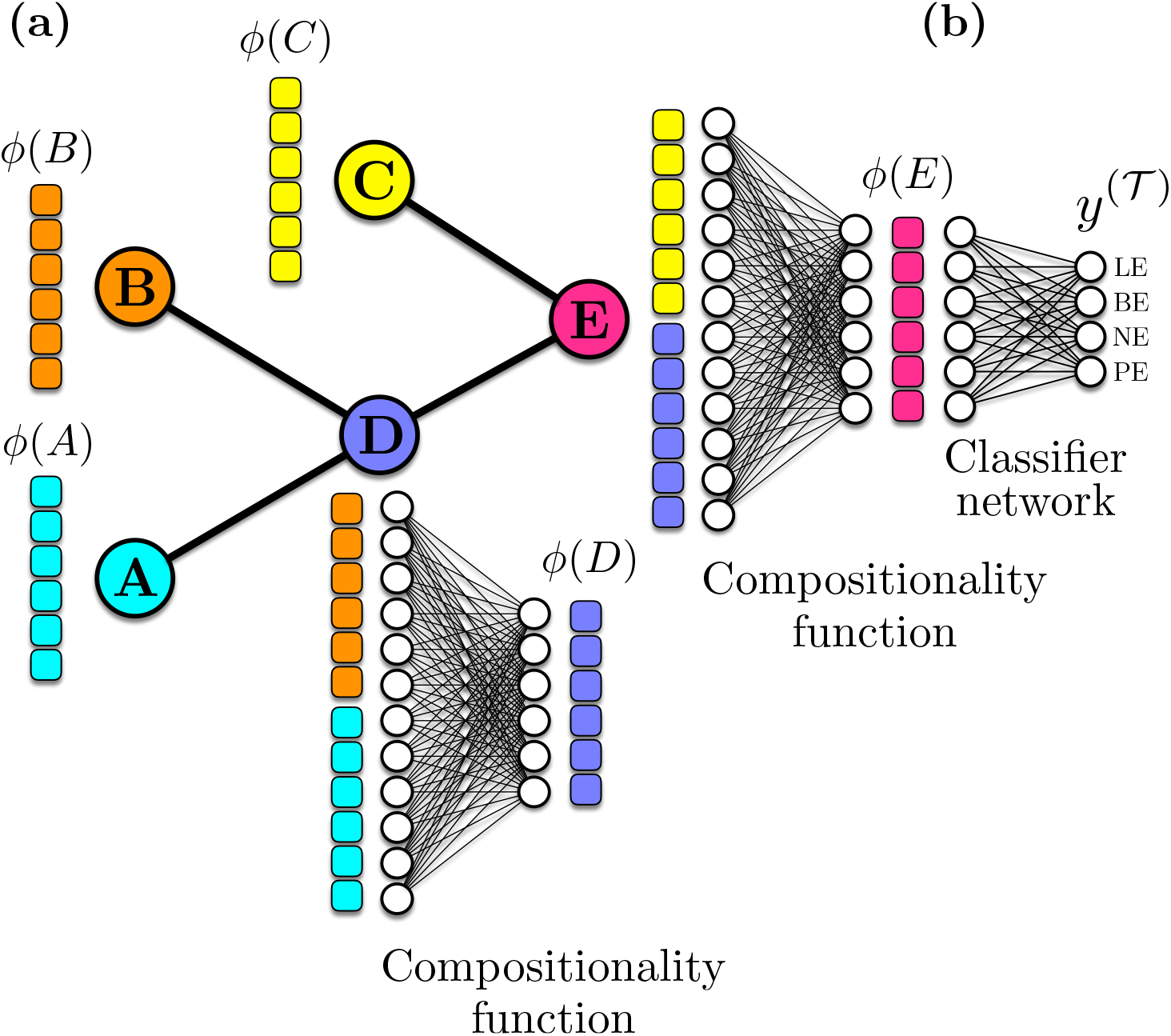
An illustrative example of processing a tree by MoTERNN. (a) Given a phylogeny and the embeddings of the leaves, MoTERNN computes the embeddings of the internal nodes up to the root of the tree in a bottom up fashion. A node and its embedding are both shown in the same color. The embeddings of *A* and *B* are given to compositionality function to produce the embedding of *D*, *ϕ*(*D*). Next, the embedding of root node, *E*, is generated from the embeddings of *C* and *D* by compositionality function (denoted by *ϕ*(*E*)). (b) The classifier network takes as input the embedding of the root node; the output contains the association scores for each mode of evolution based on which the label of the tree, 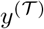 is predicted.

### 2.5 Classifier network

To predict the label/model of the phylogenetic tree, we used a classifier network. This could be a simple MLP whose weights are denoted by 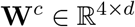. The classifier network takes as input the embedding of the root node, *ϕ*(*r*), and predicts scores for all four tumor evolutionary models, LE, BE, NE, and PE (Fig. 2b). The raw scores are passed through a softmax function to generate values between 0 and 1 that are treated as the probability values for each of the four models. The index of the maximum value corresponds the predicted label of 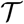 denoted by 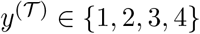. Let **z** = **W**^*c*^ · *ϕ*(*r*), formally, we have:

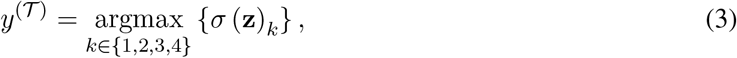

where *σ*(**z**)_*k*_ is the *k*^th^ element of the softmax activation function applied on vector **z**, defined as *σ*(**z**)_*k*_ = *e^z_k_^*/∑*_j_e^z_j_^*.

### 2.6 Loss function

We trained MoTERNN in a supervised manner using the simulated trees with their true labels (see Simulation design for more details). We used the cross-entropy loss as our objective function, which is commonly used in multi-class classification tasks. Given a tree 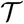 and its true label 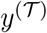 at each iteration during training, the cross-entropy between the estimated scores from the classifier network and the true labels is calculated by

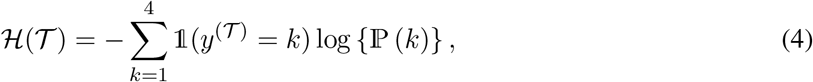

where 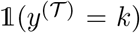 is an indicator variable that is equal to one if the true label is *k* and to zero otherwise. The second term is the logarithm of the probability of 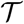 being associated with class *k*. This probability is the *k*^th^ element in the output of the softmax layer. In each iteration, 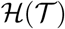 is calculated and minimized using stochastic gradient descent or its derivations [48] such as Adam [49]. The weights of the encoder network, compositionality function, and classifier network are updated through backpropagation.

### 2.7 Simulation design

The supervised learning of our model requires labeled phylogenies representing each of the four modes of cancer evolution. In general, a phylogenetic tree has two constituents, namely, a topology and branch lengths. As each mode of evolution posits conditions on the topology and branch lengths, we simulated each mode with a different scheme. To manipulate tree topologies, we used BSM, which produces binary trees with arbitrary shapes. According to BSM, the generative process of creating a tree with *N* cells includes the following steps.

1. **Sampling generative sequences**: sample a sequence of *N* – 1 independent and identically distributed (i.i.d.) random values *B* = (*b*_1_,…, *b*_*N*–1_) from the beta distribution 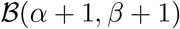, where *α* > 0 and *β* > 0 are the parameters of the beta distribution. Next, sample a sequence of i.i.d. random values *U* = (*u*_1_,…, *u*_*N*–1_) from a uniform distribution on [0, 1].
2. **Initialization**: create the root of the tree and assign the interval [0, 1] to it. Next, split the root into the left and right child nodes and assign the intervals [0, *b*_1_] and [*b*_1_, 1] to the left and right child nodes, respectively.
3. **Iteration**: in iteration *i* (considering the initialization step as the first and second iterations), among the leaves of the current tree, find the leaf whose interval [*x*, *y*] contains *u_i_*. Select the leaf and split it into the left and right child nodes. Assign [*x*, *x* + (*y* – *x*)*b_i_*] to the left child and [*x* + (*y* – *x*)*b_i_*, *y*] to the right child. Stop at iteration *N* – 1.

The parameters *α* and *β* control the shape of the tree; for example, equal values of *α* and *β* generate *balanced* topologies with high probability, and this probability increases as *α* and *β* become larger. On the other hand, the difference between the values of *α* and *β* determines the *imbalance* of the tree. We used BSM in balanced and unbalanced modes. For the balanced mode, we used (*α, β*) = (10^4^, 10^4^). A balanced topology resembles the NE mode overall topology. Also, we used it to imitate the cellular evolution within a clone. For unbalanced mode, we used (*α, β*) = (10^4^, 10^-4^). An unbalanced topology can imitate the genetic drifts as it occurs during most of a tumor’s lifetime in the LE mode.

In the following sections, we detail on our simulation schemes for the four modes of cancer evolution in terms of creating the topology and sampling the mutations on the branches of the trees.

#### 2.7.1 Simulation scheme for LE

We assume the tree topology of LE model grows during two episodes. The first and second episodes occur before and after the emergence of the dominant clone, respectively. We model the first episode with the unbalanced mode of BSM. Among the total *N* – 1 number of speciations—or cell divisions—required for simulating a tree with *N* cells, two thirds of the first speciations are done in the unbalanced mode. We model the tree growth in the second episode with the balanced mode of BSM which covers the rest of speciations. Fig. 3a shows an example of an LE tree with 20 cells. The first and second episodes are shown in orange and purple, respectively.

**Figure 3:**
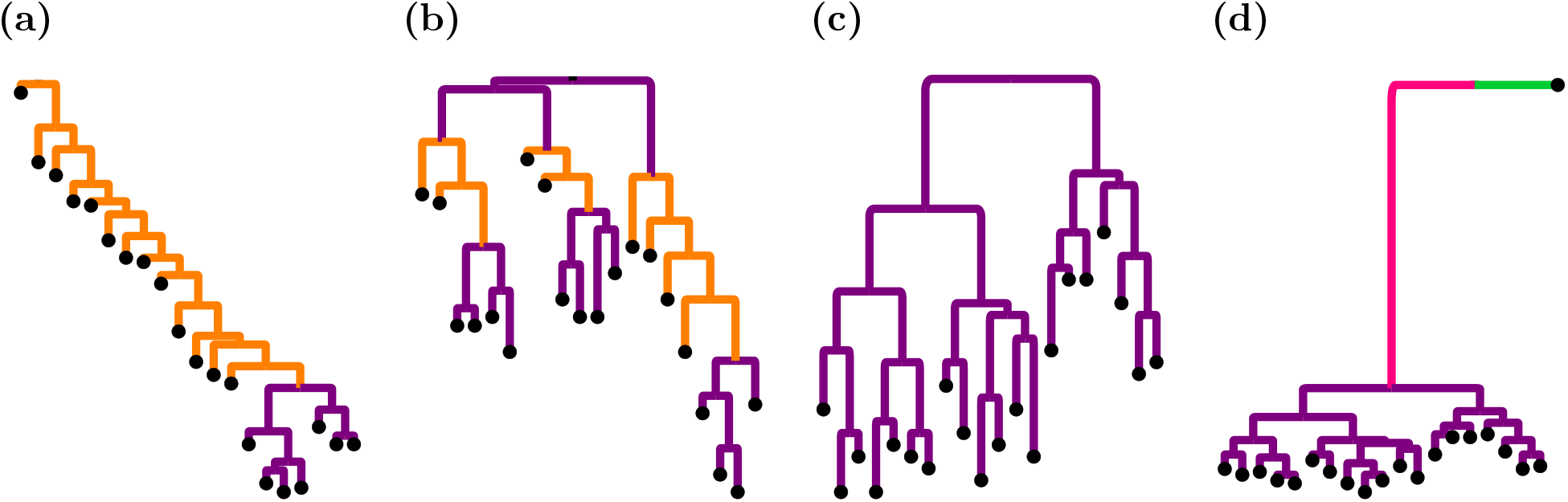
Examples of simulated trees for the four modes of cancer evolution. (a) Linear evolution. (b) Branching evolution. (c) Neutral evolution. (d) Punctuated evolution. All the trees contain 20 leaves. The branch lengths are proportional to the number of mutations. Different evolutionary episodes are shown in different colors; the branches generated using BSM’s unbalanced mode are shown in orange, while purple branches indicate balanced BSM mode of simulation. The particularly long branch in the PE mode is shown in pink. Since the input trees of MoTERNN must be binary, we added one extra branch connected to the root with no mutations in PE trees (colored in green) so that the root has in-degree 0 and out-degree 2.

After creating a topology, we sample the number of mutations from a Poisson distribution with a mean of 5 (*λ* = 5) for each branch. To generate binary genotype profiles of the cells with *M* loci, we assign an all-zeros vector to the root, which is assumed to be a normal cell without mutations. We assume the mutations are accumulated following the infinite-sites assumption (ISA) for all cancer evolution modes. Starting from the root node’s children, the tree is traversed in a breadth-first (or level-order) manner. For a branch with 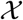 mutations, we randomly sample 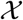 loci (without replacement) from the unmutated loci. Next, to create the genotype vector of the child node, we copy the genotype vector of the parent node and change the values of the entries corresponding to the randomly selected loci to 1.

#### 2.7.2 Simulation scheme for BE

We simulate BE trees in two steps. First, the total number of clones, *C*, is determined. We sample *C* uniformly from the set {2, 3, 4}. The tree grows using the balanced mode of BSM until C leaves are generated. In the second step, the number of cells associated with each clone is sampled. Note that the sum of the number of leaves under all clones must be equal to the total number of leaves, *N*, which is specified by the user. We sample these counts from a multinomial distribution with *N* trials and *C* categories where the success probability of each category equals 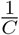. Having sampled these counts, we generate each clonal lineage using the same procedure described in Simulation scheme for LE to account for the evolutionary sweeps within each clonal lineage. Fig. 3b shows an example of a simulated BE tree.

The number of mutations on the branches and genotype profiles of the cells are generated according to the same procedure and distributions for sampling mutations in Simulation scheme for LE.

#### 2.7.3 Simulation scheme for NE

Since there are no selection or dominant clones in the NE mode, we simulate the entire tree topology using BSM’s balanced mode. The sampling of mutations and their assignment to the cells are done according to the procedure described in Simulation scheme for LE. Fig. 3c shows an example tree generated according to the NE mode.

#### 2.7.4 Simulation scheme for PE

The main characteristics of the PE trees include a long root branch with the largest portion of mutations accumulated during the lifetime of a tumor followed by a few dominant clones. To simulate such trees, we first determine the number of clones, *C*, by sampling uniformly from {2, 3}. Next, we sample the number of cells belonging to each clone from a multinomial distribution with *N* trials, *C* categories, and the success probability of 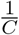 for each category (the same as in Simulation scheme for BE). Given the number of cells within each clone, we grow each clonal lineage separately using BSM’s balanced mode. The number of mutations on the long root branch is determined by sampling from a Poisson distribution with *λ* = 100. The number of mutations on the clonal branches is sampled with *λ* = 5. Since MoTERNN requires binary trees as input, we attach one extra cell with no mutations to the root that represents a normal cell. Fig. 3d shows an example tree generated according to this procedure.

## 3 Results and Discussion

### 3.1 Supervised training of MoTERNN

We simulated 8100 data points (2025 for each mode) according to the schemes described in Simulation design. A data point refers to a phylogenetic tree, a genotype matrix of the leaves, and the true label of the tree. For each phylogeny, the number of cells was sampled uniformly from the set of integers ranging from 20 to 100. We set the number of loci to 3375 to match the number of candidate loci for mutation calling in our real data set that we applied our trained model later on (see Application to real data). We applied *k*-fold cross-validation with *k* = 4 on our model to assess its predictive ability given different randomly selected test and training subsets. First, we randomly shuffled the data, and selected 100 data points as the validation set. Next, we partitioned the 8,000 remaining data points into four equal-sized subsets. In each round of cross-validation, a single subset was chosen as the test set, and the rest retained as the training set. We trained four instances of our model, separately, on each pair of training and test subsets selected by cross-validation.

We used single layer fully-connected feed-forward neural networks for all the functions, including the encoder, compositionality function, and classifier network. In particular, the encoder network was of size 3375 × 128 (3375 nodes at the input and 128 nodes at the output), the compositionality function was a network of dimensions 256 × 128 (so that it takes concatenation of two embeddings as input), and the classifier was of size 128 × 4.

In each iteration, one data point is processed to compute the cross-entropy loss. The networks’ weights are updated via the Adam optimization algorithm implemented in PyTorch. We set the learning rate of the optimizer to 10^-4^ and its weight decay parameter to zero.

In each cross-validation round, we trained MoTERNN for 6,000 iterations to process all the data points in the training set. We ran MoTERNN on an Nvidia A6000 GPU with 48 GB RAM, and an Intel Xeon Gold 6226R CPU with 64GB available RAM. The average training time was five minutes and 14 seconds. The peak of memory consumption on GPU was 1.53 GB, while on CPU, the maximum occupation of RAM was 29.6 GB.

In every iteration, we evaluated each of the four models on its entire training and validation sets. Then, we calculated the average training and validation accuracy of the four models at each iteration (Fig. 4a). The average loss function values of the models are also demonstrated in Fig. 4b. After the four models were separately trained on the cross-validation subsets, we evaluated the trained models on their corresponding test and training sets. MoTERNN achieved an average training and test accuracy of 99.98% and 99.95%, respectively.

**Figure 4:**
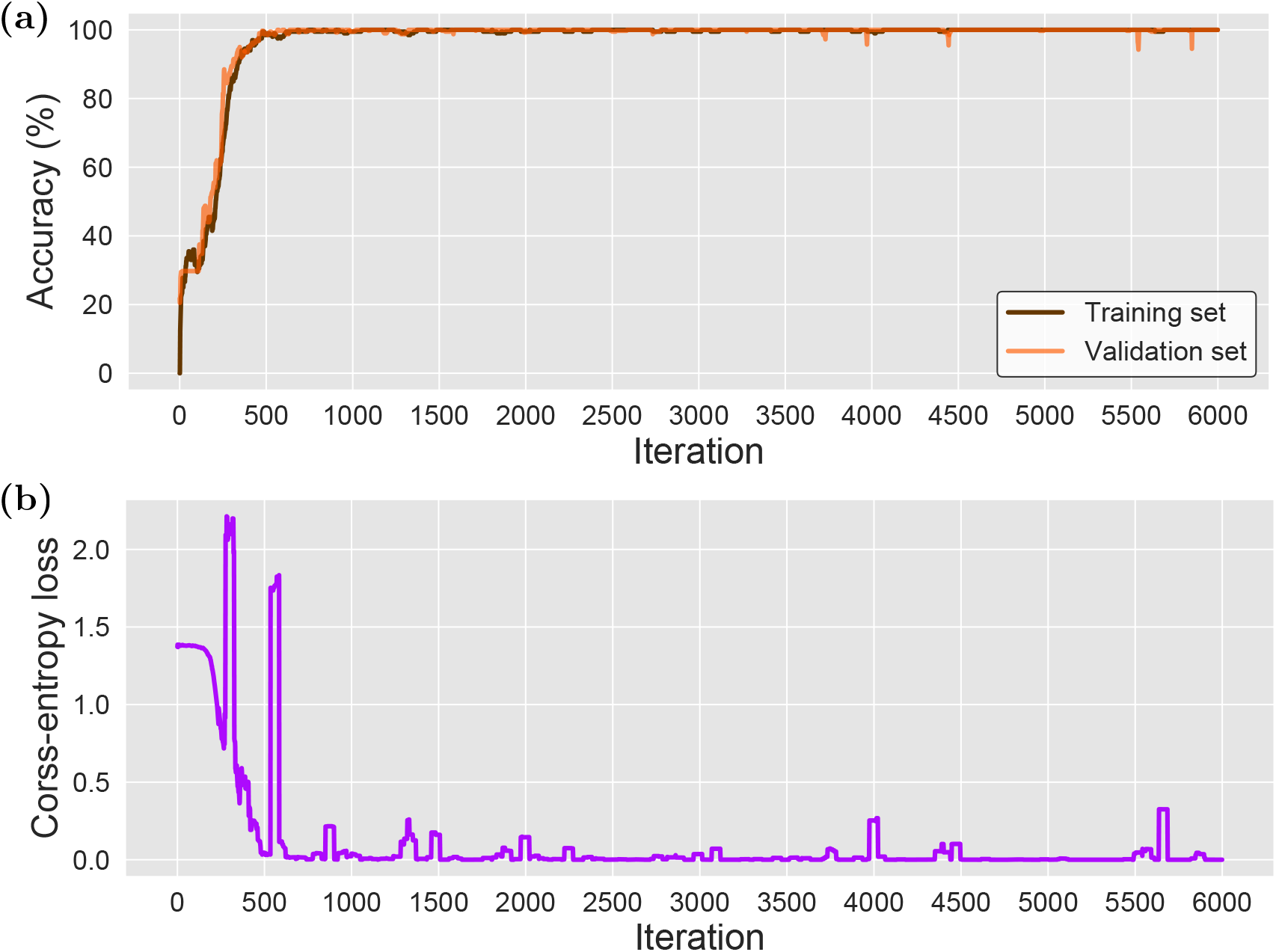
MoTERNN’s loss and classification accuracy during training. (a) The average accuracy of the four models trained in cross-validation. For each model, the accuracy on the training and validation sets was computed in each iteration. Then, we averaged over the accuracy values of the models in each iteration. (b) The average crossentropy loss function during training of the four cross-validation models. To further smooth the plots, we averaged the accuracy and loss values over the last ten iterations.

### 3.2 Application to real data

We applied our trained models on a data set consisting of single-cell whole-exome sequencing samples from a TNBC patient [45]. The TNBC data set consists of 16 diploid cells (treated as normal control samples), eight aneuploid cells, and eight hypodiploid cells [45]. We ran Phylovar [24] to infer the SNVs and the underlying phylogeny of the single-cells. The total number of candidate loci for mutation calling was 3,375. The phylogeny inferred by Phylovar is binary and admits the infinite-sites assumption [24]. To apply our trained model only on the tumor cells, we detached all the diploid cells from the phylogeny except for one that was directly connected to the root. The TNBC phylogeny is illustrated in Fig.5. Given the genotype matrix and the phylogeny, we applied the four trained models to the TNBC data and measured their average association probability scores to each mode of cancer evolution. These probability scores were approximately 0.7501, 0.2498, 0, and 0 for PE, NE, LE, and BE, respectively. Therefore MoTERNN hypothesized PE as the mode of evolution for this data set. This result is in concordance with the result of the original study. Even though [45] studied the evolutionary mode of TNBC using CNA data, and the two phylogenies—one obtained from CNA profiles and the other from SNVs—differ slightly in terms of grouping the aneuploid and hypodiploid cells [45], both analyses suggest a punctuated model of evolution for the TNBC data. It is worth mentioning that the distribution of the mutations on the branches of the TNBC’s phylogeny is different from that of the simulated data, especially, the number of mutations on the two longest branches of the TNBC phylogeny (Fig. 5) are one order of magnitude higher than the average number of mutations we assigned to the long trunk branches of PE trees in simulations (λ = 100). This implies that MoTERNN was able to recognize the overall shape of the trees regardless of the scale of the branches.

**Figure 5:**
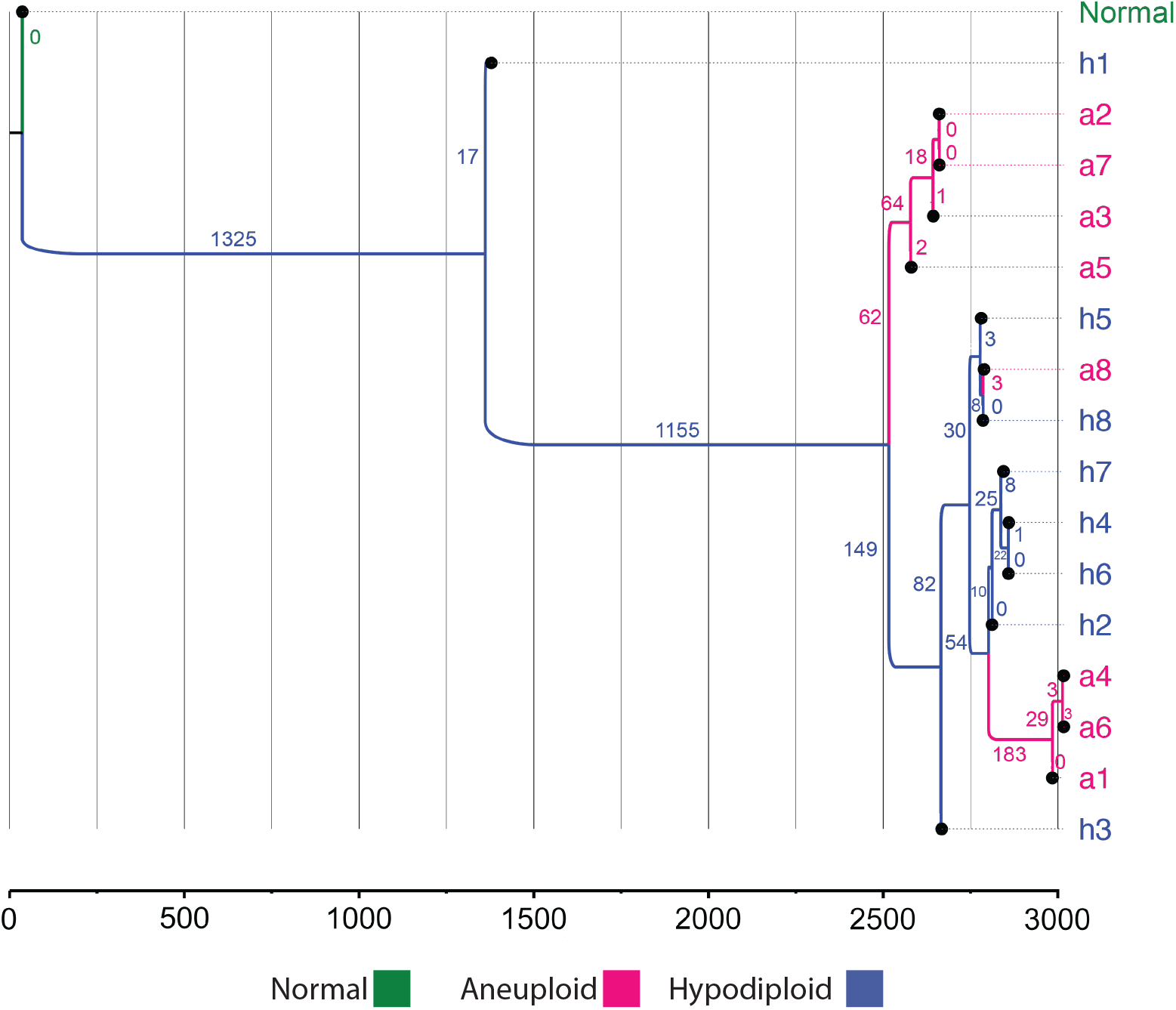
The phylogeny obtained from the TNBC data set. The aneuploid cells are tagged with *α* and colored in pink. The hypodiploid cells are tagged with *h* and colored in blue. The diploid cell connected to the root is shown in green. The branch lengths indicate the number of mutations that occurred on them. The scale axis at the bottom of the figure shows the number of accumulated mutations as the tree grows. The branches are annotated from the tips for better visualization. Figure generated by FigTree (http://tree.bio.ed.ac.uk/software/figtree/).

We note that in order to train MoTERNN for a real data set, the number of loci in the training data set must be exactly the same as in the real data set. Also, the number of cells for training must be in a range that covers the number of cells in the real data set.

## 4 Summary and Future Directions

In this work, we developed MoTERNN, a supervised learning approach to classifying the mode of cancer evolution using RNNs. MoTERNN takes as input a tumor phylogeny and the binary genotype matrix of the single-cells, and associates the tumor to one of four evolutionary modes, including LE, BE, NE, and PE. To train our model, we simulated tree topologies for the four modes of evolution and generated binary genotype matrices of the leaves following an ISA model for mutation placements on the tree branches. Our model achieved 99.98% and 99.95% accuracy on the training and test sets, respectively. We applied the trained model on a single-cell DNA sequencing data set from a triple-negative breast cancer patient. MoTERNN identified a punctuated mode of evolution in this data, in agreement with the results of the original study.

One factor that could impact the performance of our method is sampling bias. Sampling has been shown to have an impact on the inferred tree in the field of phylogenetics [50], though the debate about this has been inconclusive so far [51].

Like any other supervised learning method, the performance of MoTERNN on real data relies on the quality of simulations used to generate the training data. The ISA that we used for simulating mutations can be violated in cancer. In this regard, simulating phylogenetic trees by incorporating more complex evolutionary models such as multi-type branching evolution process [32, 35, 37] or utilizing more advanced simulators for SNVs—especially, to deviate from binary genotypes and move towards ternary genotypes— and CNAs, such as [52, 53], is a future research direction to explore.

Although learning on more advanced simulations might require more model parameters, the computational training cost would not increase dramatically by making MoTERNN more complex because such architectures train/apply, repeatedly, a few building blocks (e.g., the encoder, compositionality function, and classifier network) to the entire data structure.

As the first RNN application to phylogenetics (see [54] for details on the current status of deep learning applications in phylogenetics), MoTERNN demonstrated the potential of RNN models in learning on phylogenetic trees. We believe that variations of RNN models can be suitable choices for future studies in evolutionary biology.

## Footnotes

1 A clone consists of a group of cells with similar genotypes.

2 Sometimes the acronym RvNN is used to distinguish it from recurrent neural networks.

3 Hereafter, we use *embedding* instead of *vector embedding*.

